# Local infrared stimulation modulates spontaneous cortical slow wave dynamics in anesthetized rats

**DOI:** 10.1101/2025.09.05.674442

**Authors:** Ágnes Szabó, Richárd Fiáth, Ágoston Csaba Horváth, Péter Barthó, István Ulbert, Zoltán Fekete

**Author notes:** Corresponding author: István Ulbert, Department of Neurosurgery and Neurointervention, Faculty of Medicine, Semmelweis University, Amerikai út 57., 1145 Budapest, Hungary. These authors contributed equally to this work (Ágnes Szabó, Richárd Fiáth). Deceased, December 2024 (Zoltán Fekete).

## Abstract

Cortical slow waves are hallmark oscillations of deep sleep and certain anesthetic conditions, yet the neurobiological mechanisms controlling their dynamics remain incompletely understood. Here, we investigated the effects of local near-infrared (NIR) stimulation on slow-wave activity in ketamine/xylazine-anesthetized rats. Using a silicon-based multimodal optrode, we simultaneously delivered NIR light and recorded local field potentials (LFPs) and multi-unit activity (MUA) across cortical layers in the primary somatosensory (S1Tr) and parietal association (PtA) cortices. NIR stimulation induced local tissue heating, resulting in reproducible and reversible changes in the properties of slow waves. Specifically, up-state durations were shortened, down-states prolonged, and MUA amplitudes during up-states increased, with steeper slopes at state transitions, indicative of enhanced neuronal synchronization. LFP amplitude and spectral changes varied across cortical regions: PtA exhibited increased slow wave (0.5 – 2 Hz) and high delta (2 – 4 Hz) band activity, while S1Tr showed a trend toward reduction. Our findings demonstrate that local infrared stimulation can reliably modulate cortical slow-wave dynamics, likely through temperature-mediated changes in neuronal excitability. This approach provides a minimally invasive method for precise, local manipulation of cortical network activity and offers new insights into the biophysical regulation of slow oscillations.

## Introduction

Cortical slow waves are the hallmark of natural deep sleep, also known as slow-wave sleep (Achermann and Borbely 1997; Crunelli and Hughes 2010; Neske 2016). These high-amplitude, rhythmically recurring neural patterns not only characterize deep sleep but also appear under certain anesthetic conditions (Crunelli and Hughes 2010; Fiáth et al. 2016; Steriade et al. 1993; Ward-Flanagan et al. 2022). For instance, ketamine/xylazine anesthesia induces a stable and regular slow (∼1 Hz) rhythm composed of stereotyped slow waves that can be recorded across the entire neocortex (Fiáth et al. 2016; Sharma et al. 2010). In contrast, urethane anesthesia produces neuronal activity patterns more closely resembling those observed during natural sleep (Clement et al. 2008; Sharma et al. 2010).

At the cellular level, this slow rhythm manifests as the so-called slow oscillation, characterized by the rhythmic alternation of virtually all cortical neurons between two markedly distinct states: active periods (up-states) followed by silent periods (down-states). During up-states, the membrane potential of neurons is depolarized, resulting in elevated synaptic activity and firing rates, while during down-states, cells display a hyperpolarized membrane potential and completely suppressed spiking activity (Crunelli and Hughes 2010; Neske 2016).

Based on our current understanding, the slow oscillation is generated within the thalamocortical network, with the neocortex playing a central role (Crunelli and Hughes 2010; Timofeev et al. 2000; Sanchez-Vives and McCormick 2000). In both in vitro and in vivo models, slow waves typically originate in the infragranular layers of the cortex (particularly layer 5) and propagate in both vertical (across layers) and horizontal (across cortical regions) directions (Beltramo et al. 2013; Chauvette et al. 2010; Fiáth et al. 2016; Lőrincz et al. 2015; Sakata and Harris 2009; Sanchez-Vives and McCormick 2000; Wester and Contreras 2012). Although slow waves have a complex propagation dynamic and can initiate from virtually any cortical site, they most frequently start in frontal regions and travel toward posterior areas (Dasilva et al. 2021; Greenberg and Dickson 2013; Hangya et al. 2011; Massimini et al. 2004; Sheroziya and Timofeev 2014). Additionally, propagation of up-states has also been observed across thalamic nuclei (Horváth et al. 2024b).

Slow-wave activity is closely associated with the consolidation of memories acquired during wakefulness and with the homeostatic regulation of synaptic strengths in cortical networks (Born et al. 2006; Tononi and Cirelli 2006, 2014; Rasch and Born 2013; Timofeev and Chauvette 2017). Furthermore, previous findings suggest that slow waves contribute to the restorative function of sleep by facilitating the clearance of metabolic waste products from the brain accumulated during wakefulness (Hablitz et al. 2019; Xie et al. 2013).

Previous studies have employed various techniques to perturb the spontaneous slow rhythm in order to better understand its properties, underlying mechanisms, and functional significance. Perturbation methods resulting in rapid and transient changes of cortical activity, such as sensory, electrical, magnetic, or optogenetic stimulation, can reliably alter the ongoing brain state, for instance, by evoking or terminating up-states (Beltramo et al. 2013; D’Andola et al. 2018; Hasenstaub et al. 2007; Kuki et al. 2013; Massimini et al. 2007; Shu et al. 2003; Stoh et al. 2013; Vyazovskiy et al. 2009).

Other studies assessed the slower and more complex effects of cortical temperature on slow wave dynamics, revealing temperature-dependent changes in both spectral and temporal properties of the slow oscillation (Kalmbach and Waters 2012; Reig et al. 2010; Sheroziya and Timofeev 2015). These manipulations, however, typically involve warming or cooling relatively large cortical regions, spanning several millimeters.

Given that most slow waves emerging during non-rapid eye movement (NREM) sleep are spatially localized (Nir et al. 2011; Siclari et al. 2017), and that local slow waves also occur during rapid eye movement (REM) sleep and wakefulness (Funk et al. 2016; Siclari et al. 2017; Vyazovskiy et al. 2011), a more local manipulation of cortical temperature (and with that the indirect modulation of neuronal activity) may uncover additional, previously unrecognized aspects of slow-wave dynamics. One such technique is infrared neuromodulation - a form of photobiomodulation in which pulsed or continuous-wave infrared light is delivered into the brain - producing controlled warming in a spatially confined region of the neural tissue (Fekete et al. 2020). This photothermal effect can modulate network activity by influencing neural oscillations (Zomorrodi et al. 2019; Wang et al. 2019), and by altering the firing rates of neurons, either increasing or decreasing their excitability (Balogh-Lantos et al. 2024; Cayce et al. 2011, 2014; Horváth et al. 2020, 2022).

In this study, we applied near-infrared (NIR) stimulation to the neocortex of anesthetized rats using a penetrating multimodal probe to investigate how local temperature elevation of the tissue modulates the spectral, temporal and laminar characteristics of slow-wave activity induced by ketamine/xylazine. Cortical electrical activity was recorded simultaneously with NIR stimulation using the same device. The implanted silicon shaft of the probe served as a waveguide, focusing the emitted NIR light in the tissue near the tip of the shaft. Twelve microelectrodes, evenly spaced along the same shaft, were used to record local field potentials (LFPs) and multi-unit activity (MUA) across cortical layers. We analyzed stimulation-induced changes in the duration of up- and down-states, the amplitude of MUA during up-states, the amplitude of slow waves and the amplitude spectrum of low-frequency components (< 4 Hz) of the LFP. These features were compared between supragranular and infragranular layers, as well as between two cortical regions, namely the primary somatosensory cortex and the parietal association cortex.

## Methods

### Silicon-based optical probe

The multimodal probe (optrode) used for near-infrared stimulation and electrophysiological recordings consists of a shaft made from p-type single-crystalline silicon (Horváth et al. 2018, 2020, 2022). The shaft has a length of 5 mm and a cross-sectional area of 170 µm × 200 µm, and functions both as a waveguide and a linear electrode array. To reduce tissue damage during insertion, the shaft is tapered with a tip angle of 2α = 30°. NIR light is emitted from this sharp tip. The shaft includes 12 square-shaped recording sites (30 µm × 30 µm) arranged linearly with an inter-site spacing of 100 µm, covering a vertical area of approximately 1.1 mm. The first site is positioned 200 µm from the tip. The electrode wiring is composed of platinum (Pt), and the recording sites are coated with an additional porous Pt layer to reduce their electrical impedance to 89.6 ± 43.1 kΩ measured at 1 kHz (Horváth et al. 2022).

### Animal surgery and optrode implantation

All experiments were conducted in accordance with the EC Council Directive of September 22, 2010 (2010/63/EU), and all procedures were reviewed and approved by the Animal Care Committee of the HUN-REN Research Centre for Natural Sciences and the National Food Chain Safety Office of Hungary (license number: PE/EA/672-6/2021).

For acute in vivo experiments, adult Wistar rats were used (n = 8; weight: 331.25 ± 84.35 g, mean ± standard deviation; gender balanced). Surgical procedures, probe implantation, and electrophysiological recordings were performed similarly to our previous studies (Balogh-Lantos et al. 2024; Fiáth et al. 2016; Horváth et al. 2024a; Ismaiel et al. 2023). Briefly, anesthesia was induced via intraperitoneal injection of ketamine (75 mg/kg) and xylazine (10 mg/kg). To maintain a stable and regular cortical slow oscillation throughout the experiment, supplementary doses of ketamine/xylazine were administered intramuscularly at hourly intervals. A homeothermic heating pad connected to a temperature controller (Supertech, Pécs, Hungary) was used to maintain physiological body temperature. The head of the anesthetized rat was secured in a stereotaxic frame (David Kopf Instruments, Tujunga, CA, USA), after which the skin and the connective tissue were removed from the top of the skull.

In four animals, a cranial window measuring approximately 10 mm × 5 mm was drilled over the left brain hemisphere, centered around the parietal association cortex (PtA; anterior-posterior [AP]: +1 mm to −9 mm; medial-lateral [ML]: 0.5 mm to 5.5 mm; coordinates relative to bregma (Paxinos and Watson 2007). This large craniotomy was required to accommodate a flexible micro-electrocorticography (µECoG) array placed on the exposed cortical surface (Csernyus et al. 2021; Ismaiel et al. 2023). This device was used to collect electrophysiological data for a separate study. The 32-channel µECoG array featured a circular opening (800 μm diameter) through which the optrode was implanted for infrared stimulation and intracortical recording. In the remaining four animals, a smaller craniotomy with a size of 3 mm × 3 mm was prepared (AP: −1.5 mm to −4.5 mm; ML: 1.5 mm to 4.5 mm, with respect to bregma), targeting the trunk region of the primary somatosensory cortex (S1Tr) for optrode insertion.

At the insertion site, the dura mater was carefully removed using a 34-gauge needle to minimize cortical dimpling during optrode implantation. The optrode was mounted on a motorized stereotaxic micromanipulator (Robot Stereotaxic, Neurostar, Tübingen, Germany) and inserted perpendicularly to cortical layers to a depth of 1.2 mm with a slow rate (2-20 μm/s) to reduce the insertion-related mechanical damage to the tissue (Fiáth et al. 2019). After reaching the target depth, the tip of the optrode was located in infragranular cortical layers (bottom of layer 5, or layer 6; Fig 1A). The tip position of optrode was chosen based on previous findings indicating a key role for deep layers in the initiation and maintenance of up-states (Beltramo et al. 2013; Chauvette et al. 2010; Fiáth et al. 2016; Lőrincz et al. 2015; Sakata and Harris 2009; Sanchez-Vives and McCormick 2000; Wester and Contreras 2012). To prevent dehydration of the cortical tissue, room temperature physiological saline solution was regularly applied to the brain surface.

**Figure 1.**
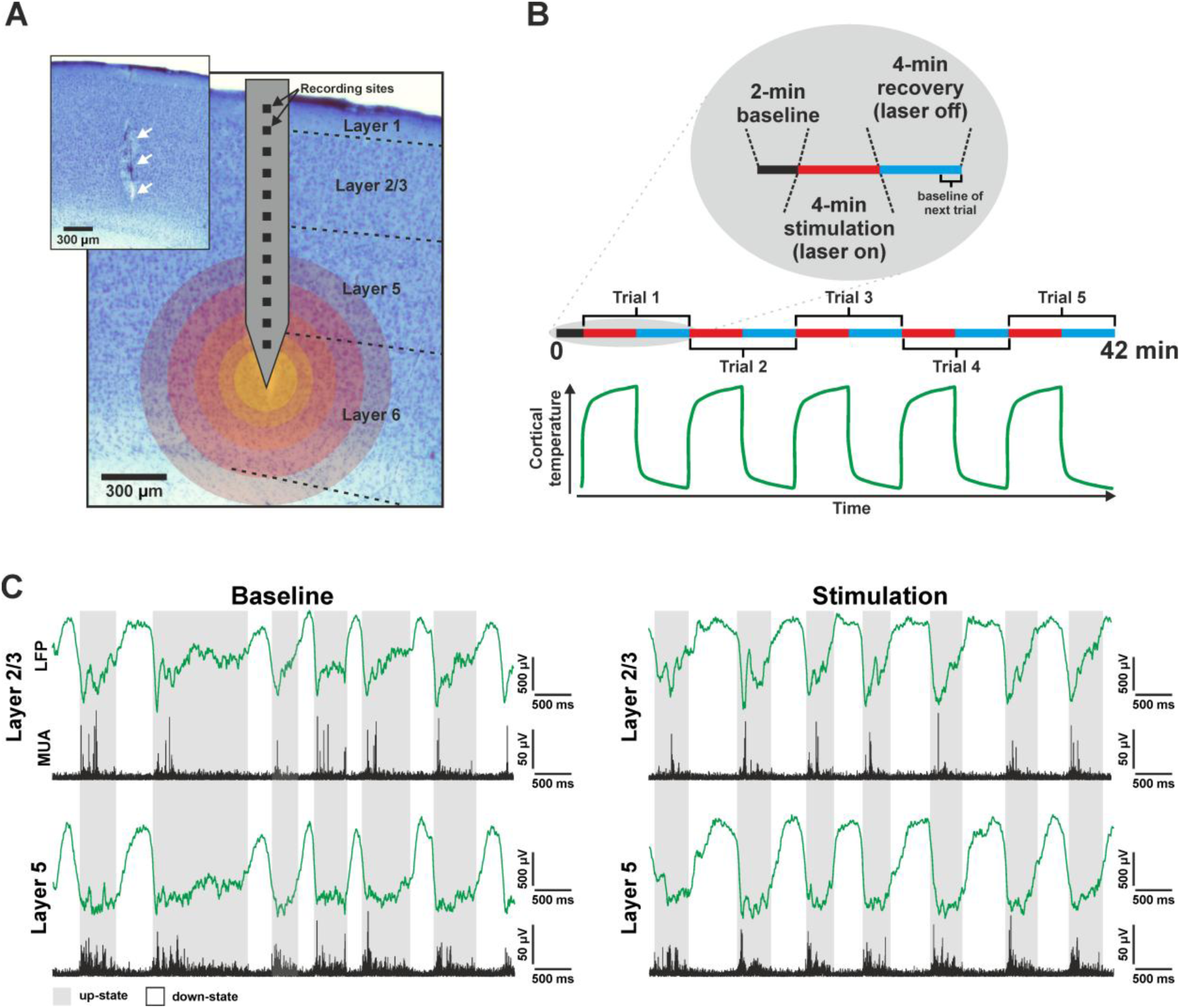
(A) Experimental setup. Schematic illustration of the silicon-based optical probe (optrode) overlaid on a Nissl-stained coronal brain section containing the optrode track (indicated by white arrows in the inset). Boundaries of cortical layers are marked with dashed black lines. Colored circles overlaid on the image indicate part of the cortical area affected by infrared stimulation. Lighter colors closer to the optrode tip indicate a higher increase in cortical temperature during stimulation. (B) Schematic illustration of the infrared stimulation protocol. In 2 out of the 8 rats, stimulation was applied only for 2 minutes. Bottom: schematic plot showing the time course of cortical temperature during the stimulation protocol. (C) Representative five-second local field potential (green) and multi-unit activity (black) traces recorded from layer 2/3 (top) and layer 5 (bottom) of the neocortex during the baseline period (last minute of an OFF period, left) and during infrared stimulation (last minute of an ON period, right) from a single rat.

### In vivo electrophysiological recordings

Cortical electrical activity was collected with the optrode using an Intan RHD2000 electrophysiological recording system (Intan Technologies, Los Angeles, CA, USA) equipped with a 32-channel headstage. Wideband (0.1 – 7500 Hz) continuous signals were acquired at a sampling rate of 20 kHz per channel (30 kHz per channel for one animal) with 16-bit resolution. For the four animals with the µECoG array, a second 32-channel headstage was used to record ECoG signals; however, this data was not analyzed in the present study. A stainless-steel needle inserted in the nuchal muscle of the animal served as both the reference and ground electrode during recordings.

### Infrared neuromodulation

NIR light at a wavelength of 1550 nm was delivered using a fiber-coupled laser diode with a maximum output power of 70 mW (LPSC-1550-FG105LCA-SMA, Thorlabs GmbH, Bergkirchen, Germany). The laser was driven at a constant current of 400 mA DC, supplied by a Keithley 2611B SYSTEM SourceMeter (Keithley Instruments Inc, Cleveland, OH, USA). Two slightly different stimulation protocols were applied during the experiments. Following a 2-minute-long baseline recording period, the laser diode was activated for either 2 minutes (n = 2 rats) or 4 minutes (n = 6 rats), referred to as ON period. NIR stimulation resulted in an estimated temperature increase of approximately 4.5 °C and 4.8 °C for 2-min and 4-min ON periods, respectively (measured in vitro; Balogh-Lantos et al. 2024). Each ON period was followed by a 4-minute OFF period, during which the laser diode was turned off, allowing the tissue and the temperature to recover (Fig. 1B). This ON-OFF cycle was repeated five times consecutively, resulting in recordings with a duration of either 32 minutes (n = 2 rats) or 42 minutes (n = 6 rats). Repetitions will be referred to as trials. The onset of ON and OFF periods was synchronized with the electrophysiological recordings using trigger signals generated by the same Keithley instrument.

### Histology

To visualize the track of the optrode in the brain tissue and to identify cortical layers (Fig. 1A), we followed a histological procedure described previously (Fiáth et al. 2016; Fiáth et al. 2019). In short, a subset of rats (n = 5) was deeply anesthetized with ketamine/xylazine after neural data collection and transcardially perfused with 100 ml physiological saline solution followed by 250 ml of a fixative solution containing 4% paraformaldehyde in 0.1 m phosphate buffer (pH 7.4). The fixed brains were stored at 4 °C overnight. Subsequently, the brain tissue was cut into 60-μm-thick coronal sections using a vibratome (Leica VT1200, Leica Microsystems, Wetzlar, Germany). After washing in 0.1 M phosphate buffer, the sections were transferred to a Petri dish containing gelatin, mounted onto microscopic slides, and air-dried. The slides were then processed for cresyl violet (Nissl) staining, dehydrated in xylene, and cover-slipped using DePex (SERVA Electrophoresis, Heidelberg, Germany). Finally, Nissl-stained sections containing the optrode tracks were photographed under a light microscope (Leica DM2500, Leica Microsystems) equipped with a digital camera (DP73, Olympus, Tokyo, Japan). The track of the optrode was clearly visible in the brain sections due to the relatively large cross-sectional area of its silicon shaft.

### Data analysis

All electrophysiological data were processed offline with MATLAB (MathWorks, Natick, MA, USA)-based algorithms. Since cortical temperature returns to baseline values within two minutes after cessation of NIR stimulation (Ismaiel et al. 2023), we defined the last minute of the four-minute-long OFF period as the baseline (BL) period for the subsequent stimulation trial (Fig. 1B). For the first stimulation trial, the baseline period was the second minute of the cortical recording containing spontaneous activity. All subsequent analyses were applied separately for each minute of the ON period. In most cases, however, we compared the values calculated during the baseline period with those computed during the last minute of the stimulation period, during which the effect of infrared stimulation was strongest. Computed values were averaged across the five trials and across channels corresponding to the same layer.

#### Laminar location of recording sites

The tip of the probe was positioned in the infragranular cortical layers, while microelectrodes recorded simultaneously from multiple layers. To determine the precise laminar location of optrode sites, we utilized both the registered cortical activity patterns and, where available, anatomical information (Nissl-stained coronal brain sections containing the optrode track). Each recording site was assigned to a specific cortical layer, based on criteria established in our previous work (Fiáth et al. 2016).

Under ketamine/xylazine anesthesia, superficial layers typically exhibit sparse spontaneous population activity, with fewer neurons active compared to deeper layers (Barth et al. 2012; Fiáth et al. 2016; Sakata and Harris 2009). Due to the absence of spiking activity in layer 1, sites localized to this layer were excluded from further analysis. Recording sites located within layers 2 and 3 were grouped together. Since layer 4 is a thin layer and virtually absent in the parietal association cortex, data from sites potentially located in this layer (in S1Tr) were also excluded.

Layer 5, which contains the cell bodies of the largest pyramidal cells across layers, displays the strongest neuronal activity under ketamine/xylazine anesthesia (Fiáth et al. 2016). This layer also plays a key role in the generation of slow waves (Chauvette et al. 2010; Fiáth et al. 2016; Lőrincz et al. 2015; Sanchez-Vives and McCormick 2000). Recording sites in layer 5 were typically used as a reference for identifying other layers, as this layer can be located with the greatest reliability. Finally, since only relatively few sites were located in layer 6, activity from this layer was not analyzed. Thus, the present study focused primarily on neuronal activity in two cortical layers: layer 2/3 and layer 5.

#### Multi-unit activity-based state detection

To examine NIR stimulation-related changes in the duration of up- and down-states and in neuronal population activity (multi-unit activity; MUA, Fig. 2), we first identified the onset of slow oscillation states. The MUA-based state detection algorithm applied here was adapted from Fiáth et al. (2016). Briefly, MUA was extracted by bandpass filtering the wideband recordings between 500 - 5000 Hz with a third-order, zero-phase shift Butterworth filter. The resulting signal was then rectified and downsampled from 20 kHz to 2 kHz. Downsampled MUA signals from all twelve channels were summed (excluding nonfunctional or noisy channels), and the resulting single-channel time series was then smoothed to obtain the envelope of cortical population activity with a third-order, zero-phase shift Butterworth lowpass filter (50 Hz cutoff frequency). This signal was termed as smoothed population activity (SPA). Finally, to detect up- and down-state onset times, an amplitude threshold was computed based on the mean and standard deviation of short (50 ms) periods of the SPA signal selected within down-states. The detection algorithm imposed two constraints: a minimum up-state duration of 50 ms and a minimum down-state duration of 100 ms. The detected up- and down-state onsets were saved for subsequent analysis.

**Figure 2.**
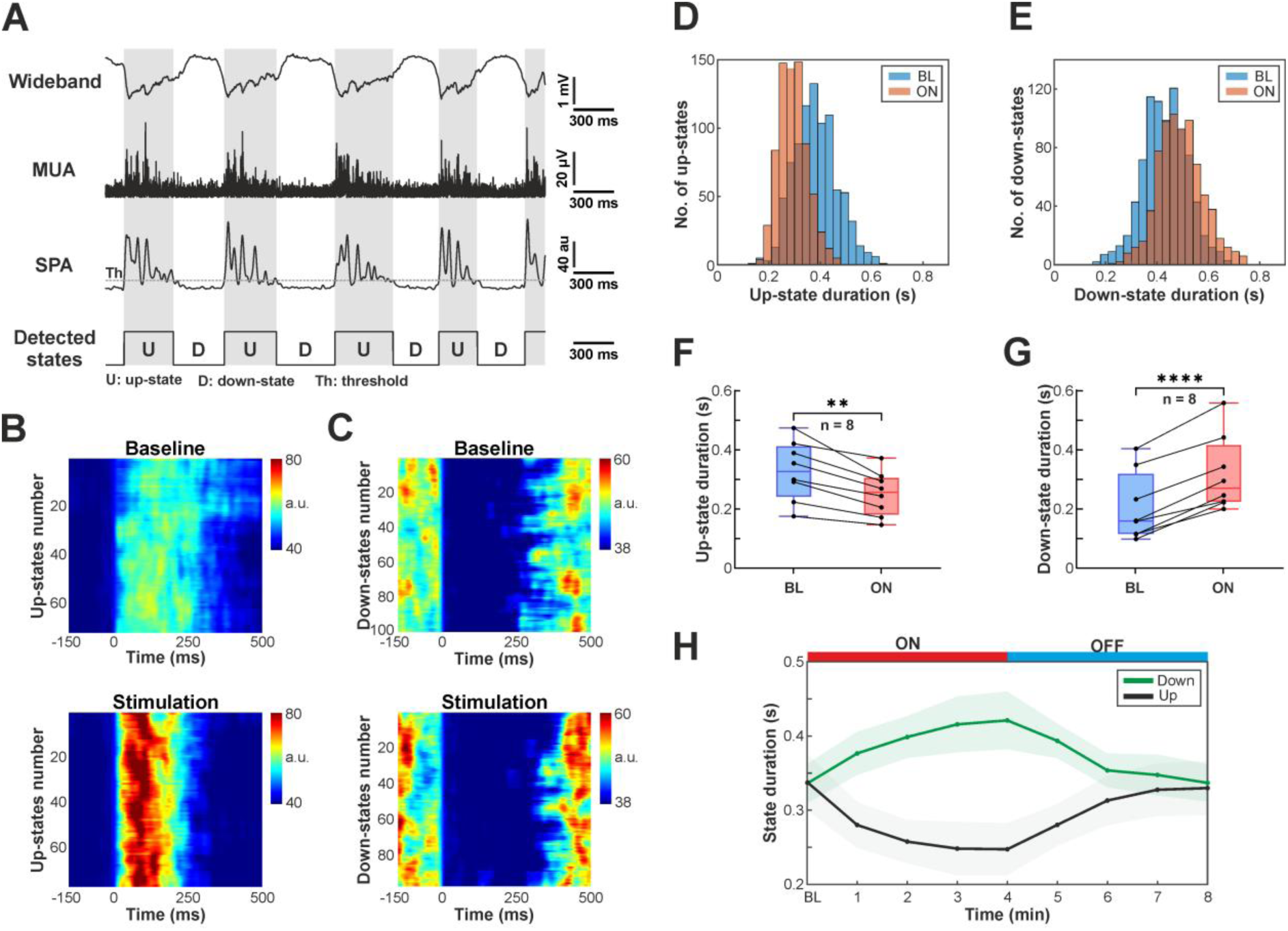
(A) Schematic illustration demonstrating the method used for detection of up- and down-state onsets. MUA, multi-unit activity; SPA, smoothed population activity. (B-C) Color maps of short segments of MUA traces aligned to the detected up-state (B) and down-state (C) onsets (time point zero), ordered by occurrence. The color maps display SPA traces extracted during the last minute of the OFF period (baseline, top) and the last minute of the ON period (stimulation, bottom) of a single trial. (D-E) Representative distributions of up-state (D) and down-state (E) durations during baseline (last two minutes of OFF periods; blue) and stimulation (last two minutes of ON periods; red) periods. (F-G) Distribution of mean up-state (F) and down-state (G) durations across all rats during baseline (BL) and stimulation (ON) periods. **, p < 0.01; ****, p < 0.0001. (H) Changes in mean up-state (black) and down-state (green) durations during stimulation trials. The red and blue horizontal bars indicate the stimulation (ON) and recovery (OFF) periods, respectively. The shaded areas indicate standard error.

#### Calculation of up- and down-state durations

Up- and down-state durations were computed using the previously extracted up- and down-state onset times. The duration of a given up-state (or down-state) was defined as the time interval between the onset of the current up-state (or down-state) and the onset of the following down-state (or up-state). To improve reliability, up- and down-states with excessively short or long durations (i.e., exceeding three scaled median absolute deviations from the median), which likely resulted from inaccurate detections, were excluded from the analysis.

#### Computation of up-state onset-locked multi-unit activity averages

To calculate MUA averages aligned to the start of up-states for both the stimulation (ON) and the recovery (OFF) period (Fig. 3), we adapted the method described by Horváth et al. (2024a). Briefly, short segments were extracted from the continuous MUA recordings around each detected up-state onset (from 150 ms before to 500 ms after the onset). For each stimulation trial (separately for ON and OFF periods), these segments were then averaged across up-states and across cortical channels corresponding to either layer 2/3 or to layer 5. Resulting segments were subsequently averaged across trials. Only up-states detected during the last minute of the ON (471.67 ± 118.93 up-states per animal; mean ± standard deviation) and OFF periods (468.63 ± 111.17 up-states per animal) were used to compute the MUA averages. Finally, we calculated the average MUA level within a 40 ms window starting 10 ms after the up-state onset, both during stimulation and the baseline period.

**Figure 3.**
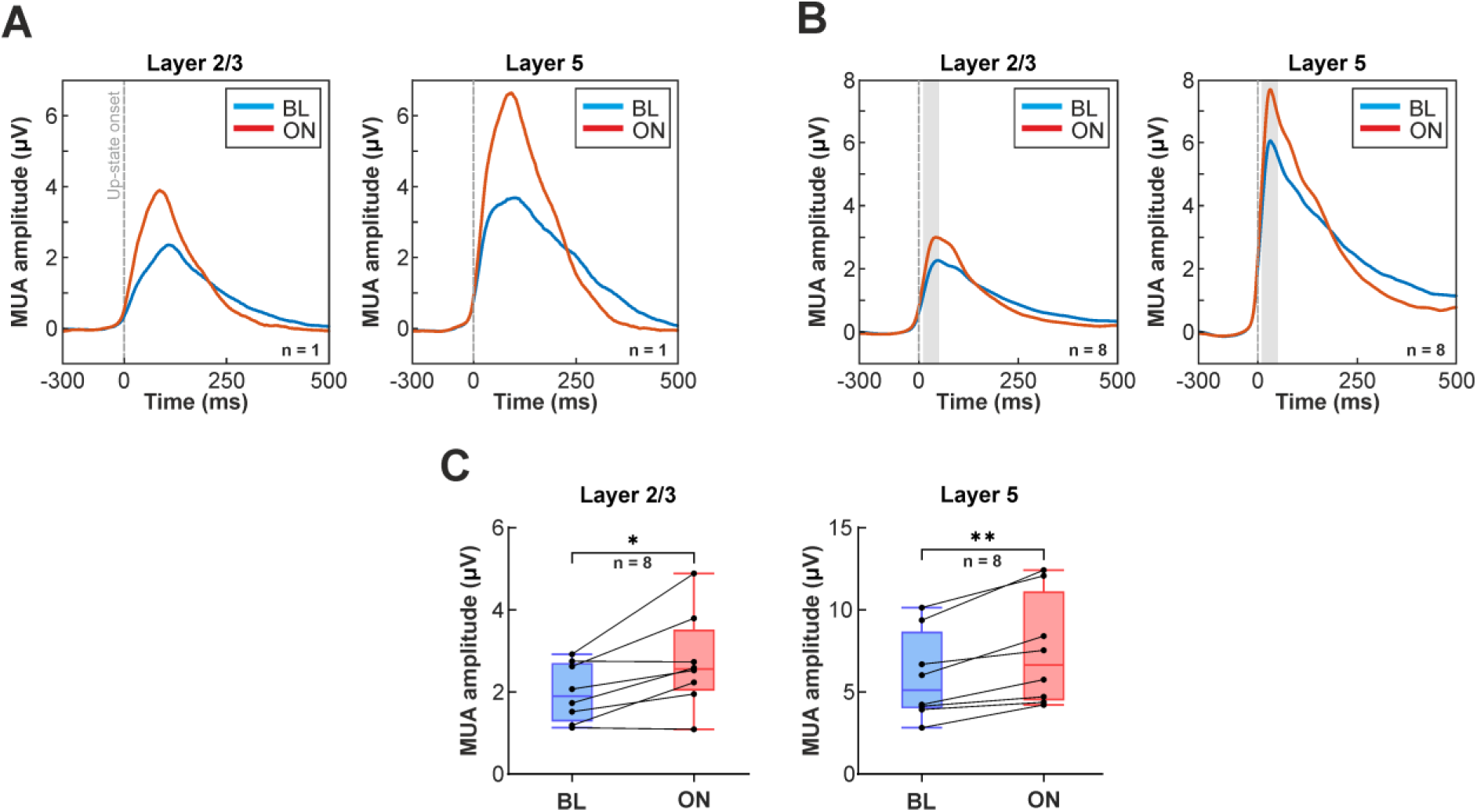
(A) Representative examples of multi-unit activity (MUA) traces averaged across up-states from one rat during baseline (BL, blue) and infrared stimulation (ON, red) in layer 2/3 (left) and layer 5 (right). The up-state starts at time point zero (dashed gray vertical line). (B) Average of up-state onset locked MUA across all rats (n = 8) during baseline (blue) and stimulation (red) in layer 2/3 (left) and layer 5 (right). The shaded gray area indicates the time window used to calculate MUA amplitude values shown in panel C. (C) Distribution of mean MUA amplitudes calculated within a 40-ms time window (10-50 ms after up-state onset; see panel B) during baseline and stimulation in layer 2/3 (left) and layer 5 (right). *, p < 0.05; **, p < 0.01.

#### Comparison of local field potential amplitudes

We also investigated whether NIR stimulation had a measurable effect on the amplitude of cortical slow waves (Fig. 4). To extract local field potentials, the recorded wideband (0.1 - 7500 Hz) cortical signals were bandpass filtered between 0.5-200 Hz with a third-order, zero-phase shift Butterworth filter. To eliminate the power line noise, a second-order infinite impulse response notch filter was applied at 50 Hz. To reduce data size and computation time, LFP signals were downsampled to 2 kHz. Nonfunctional and noisy channels were removed from further analysis (n = 8 channels in total across the 8 recordings).

**Figure 4.**
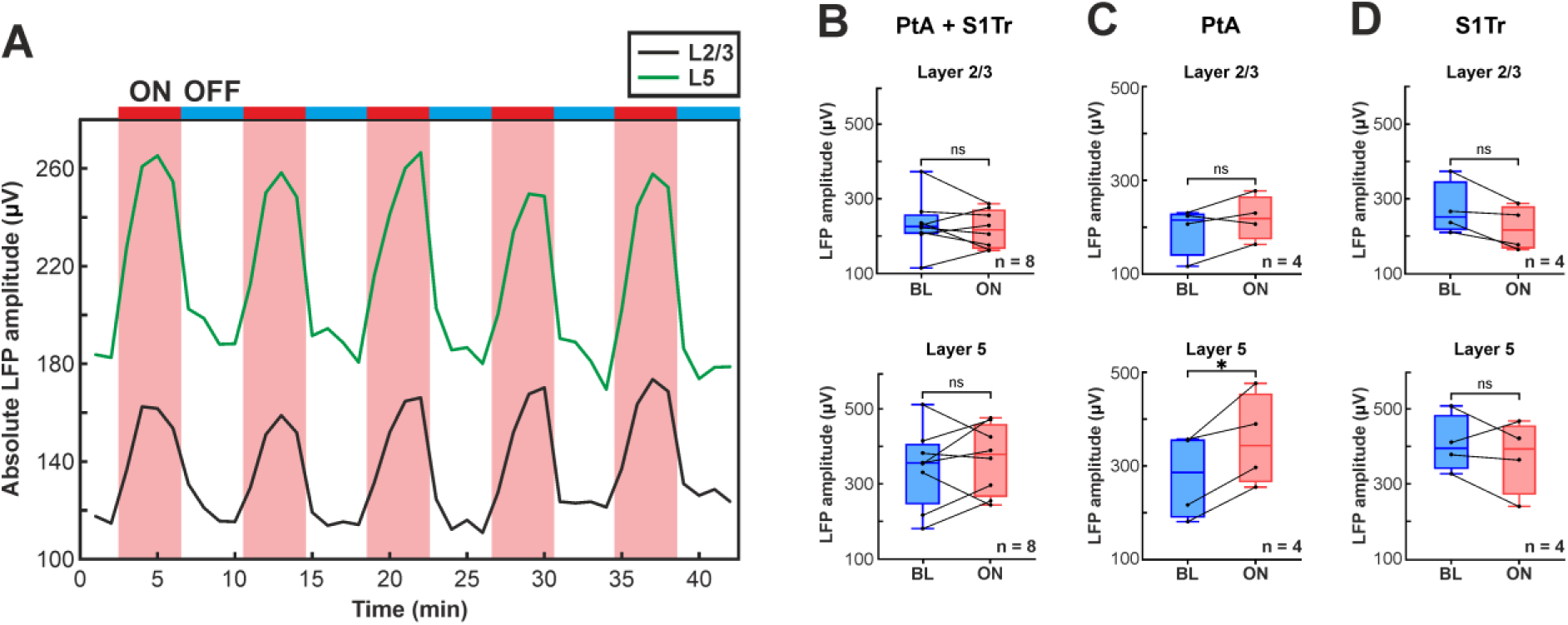
(A) Representative example of changes in the absolute LFP amplitude over time in layer 2/3 (black) and layer 5 (green) during the infrared stimulation protocol. The red and blue horizontal bars indicate the stimulation (ON) and recovery (OFF) periods, respectively. (B) Distribution of mean absolute LFP amplitudes during stimulation (ON) compared to baseline (BL) periods in all rats in layer 2/3 (top) and layer 5 (bottom). (C) Same as panel B, but restricted to PtA data. (D) Same as panel B but, restricted to S1Tr data. *, p < 0.05; ns, not significant. PtA, parietal association cortex; S1Tr, trunk region of the primary somatosensory cortex.

To assess the influence of NIR stimulation on LFP amplitudes, we calculated and compared the mean absolute LFP amplitude during the last minute of ON and OFF periods, the latter used as baseline. Amplitudes were also averaged across trials and across channels corresponding to the same layer.

#### Spectral analysis of local field potentials

Fast Fourier transformation (FFT) was applied to the extracted LFP signal to calculate the amplitude spectrum in the low-frequency band from 0.5 Hz to 4 Hz (Fig. 5B). Next, we calculated the average spectral amplitude in two separate bands: from 0.5 Hz to 2 Hz, referred to as the slow wave band, and from 2 Hz to 4 Hz, referred to as high delta band. The FFT amplitude spectrum was computed separately for the last minute of the ON and OFF periods, as well as for each channel. Spectral amplitude values were averaged across trials and across channels corresponding to the same cortical layer. To visualize spectral changes during the entire recording, spectrograms were computed using the Chronux MATLAB package (Bokil et al. 2010; Fig. 5A).

**Figure 5.**
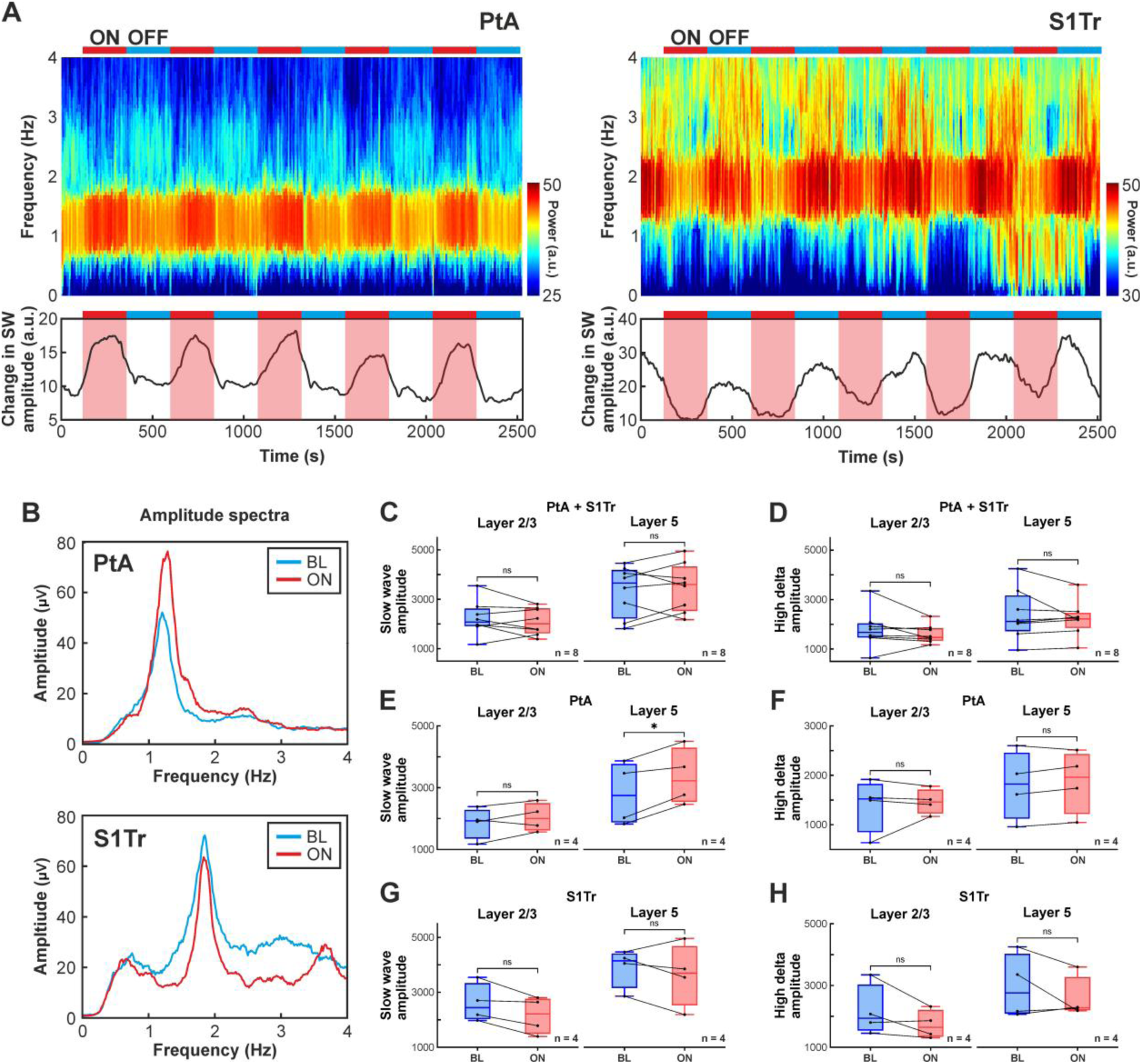
(A) Spectrograms computed from layer 5 local field potentials demonstrating spectral changes during the whole infrared stimulation protocol in two rats. In one rat (optrode implanted in PtA, left), the power of low-frequency (0.5-4 Hz) activity increased during stimulation, while in the other rat (optrode implanted in S1Tr, right), a decrease in spectral power occurred during stimulation. The change in the power of the slow wave band (0.5 – 2 Hz) is shown below the spectrograms. The red and blue horizontal bars indicate the stimulation (ON) and recovery (OFF) periods, respectively. (B) Representative FFT amplitude spectra from layer 5 of the same two rats in the low-frequency band (0.5 - 4 Hz) during baseline (blue) and stimulation (red) periods. (C-D) Distribution of mean spectral amplitudes in the slow wave (0.5 - 2 Hz; C) and high delta (2 – 4 Hz, D) bands during stimulation (ON) compared to baseline (BL) in all rats in layer 2/3 (left) and layer 5 (right). (E-F) Same as panels C-D but restricted to PtA data. (G-H) Same as panels C-D but restricted to S1Tr data. *, p < 0.05; ns, not significant. PtA, parietal association cortex; S1Tr, trunk region of the primary somatosensory cortex.

### Statistical analysis

All statistical analyses were performed using GraphPad (v10.3.1). Normality of the data was assessed using the Shapiro–Wilk and Kolmogorov-Smirnov tests, and Student’s t-test or paired t-test was used to compare groups. A p-value < 0.05 was considered statistically significant. Box-and-whisker plots (see Figs. 2-4) showing the distribution of data are presented as follows. The middle line indicates the median, the boxes correspond to the 25th and 75th percentiles, and the whiskers mark the minimum and maximum values. Individual values are depicted as black dots. Unless otherwise stated, all values in the Results section are reported as mean ± standard deviation.

## Results

### Infrared stimulation modulates spontaneous cortical slow-wave activity in anesthetized rats

We used a multimodal probe (referred to as an optrode) with its tip inserted into the infragranular layers of the neocortex of ketamine/xylazine anesthetized rats (n = 8) to deliver near-infrared light stimulation while simultaneously recording cortical local field potentials and multi-unit activity. The optrode contained twelve, equidistantly spaced microelectrodes, allowing for recordings across multiple cortical layers (Fig. 1A). The device was implanted either in the parietal association cortex (PtA, n = 4 rats) or in the trunk region of the primary somatosensory cortex (S1Tr, n = 4).

In each animal, continuous-wave NIR stimulation was applied for 4 minutes (ON period, 2 minutes in 2 rats), followed by a 4-minute recovery period without stimulation (OFF period; Fig. 1B). Each ON-OFF cycle was repeated five times per animal, with all recordings starting with a two-minute baseline period.

NIR stimulation locally elevates cortical temperature by several degrees Celsius, with the most pronounced increase occurring within 1 mm of the probe tip (Balogh-Lantos et al. 2024; Horváth et al. 2020, 2022). This stimulation-induced hyperthermia led to marked alterations in the properties of ketamine/xylazine-induced cortical slow-wave activity. Changes were observed in both low-frequency (< 200 Hz; LFP) and high-frequency (500-5000 Hz; MUA) components of the recorded neural signal (Fig. 1C). For instance, both the duration of up- and down-states and the amplitudes of slow waves were modulated during stimulation compared to baseline. These effects persisted throughout stimulation and were fully reversible, with parameters rapidly returning to baseline once NIR stimulation ceased. Furthermore, the effects of local cortical warming on slow wave properties were consistent and reproducible across repeated stimulation trials.

To assess potential layer-specific effects, we analyzed stimulation-induced responses separately in supragranular (i.e., layer 2/3) and infragranular (i.e., layer 5) cortical layers (Fig. 1C). In addition, we compared the effects of stimulation between cortical areas (PtA vs. S1Tr).

### Effect of infrared stimulation on the duration of up- and down-states

Cortical slow waves consist of two alternating phases: up-states, characterized by high synaptic activity and neuronal spiking, and down-states, marked by near-complete neuronal silence. Under ketamine/xylazine anesthesia, both phases typically last several hundred milliseconds. Previous studies have demonstrated that the duration of these states is sensitive to tissue temperature (Kalmbach and Waters 2012; Reig et al. 2010). We therefore examined whether NIR stimulation alters the duration of up- and down-states.

Up- and down-state onsets were detected using a method based on cortical population activity (Fig. 2A). This approach typically provides precise estimates of state onset times and allows accurate measurement of individual state durations (Fiáth et al. 2016). We found no statistically significant differences in baseline up- and down-state durations between PtA and S1Tr, although up-states were slightly longer in PtA (PtA vs. S1Tr; up-state: 362.28 ± 86.15 ms vs. 292.24 ± 114.16 ms, p = 0.308, Student’s t-test; down-state: 329.25 ± 69.95 ms vs. 340.85 ± 94.06 ms, p = 0.826, Student’s t-test).

Visualization of population activity across states detected during stimulation and recovery revealed clear changes in the duration of up- and down-states (Fig. 2B and C). The distribution of state durations also shifted noticeably during stimulation (Fig. 2D and E). Specifically, NIR stimulation significantly reduced up-state duration (BL vs. ON; 320.89 ± 94.87 ms vs. 262.40 ± 85.29 ms; p = 0.0012, paired t-test; Fig. 2F), while down-states became significantly longer relative to baseline (BL vs. ON; 337.01 ± 76.29 ms vs. 384.71 ± 81.16 ms; p < 0.0001, paired t-test; Fig. 2G). Consequently, the proportion of time spent in up-states decreased from 48.77% to 37.38%, while down-states increased from 51.23% to 62.62% during stimulation. Despite these changes, the total duration of slow oscillation cycles (the sum of the duration of a single up- and down-state) remained largely unchanged during the whole recording, which was found 662.31 ms and 660.40 ms for the last minute of OFF and ON periods, respectively (Fig. 2H). Changes in state durations progressed gradually during both stimulation and recovery (Fig. 2H).

### Effect of infrared stimulation on population activity

In addition to reducing the duration of up-states, NIR stimulation also increased MUA amplitudes (Fig. 2B). To investigate this effect further, we computed the average MUA aligned to the onset of detected up-states in supragranular (layer 2/3) and infragranular (layer 5) layers, separately for stimulation and recovery periods (Fig. 3A and B).

Consistent with previous findings (Fiáth et al. 2016), MUA amplitude was significantly higher in layer 5 compared to layer 2/3, (layer 2/3 vs. layer 5; 1.99 ± 0.66 µV vs. 5.91 ± 2.50 µV, p = 0.0024, paired t-test). No significant differences were observed between cortical regions, although MUA amplitudes tended to be slightly higher in S1Tr (PtA vs. S1Tr; layer 2/3: 1.85 ± 0.55 µV vs. 2.13 ± 0.74 µV, p = 0.6134, Student’s t-test; layer 5: 5.70 ± 2.54 µV vs. 6.13 ± 2.43 µV, p = 0.8395, Student’s t-test).

In most animals, MUA amplitude during up-states was higher during NIR stimulation compared to baseline (Fig. 3A and B). Additionally, the slope of MUA traces was steeper at both down-to-up and up-to-down transitions, indicating sharper and more synchronous state transitions during stimulation.

To quantify this effect, we measured the mean MUA amplitude within a 40-ms window (10-50 ms) following the up-state onset during the last minute of the stimulation period and compared it with the corresponding baseline period (last minute of recovery). MUA amplitudes were significantly higher during NIR stimulation in both cortical layers (BL vs. ON; layer 2/3: 1.99 ± 0.66 µV vs. 2.72 ± 1.08 µV, p = 0.0174, paired t-test; layer 5: 5.91 ± 2.50 µV vs. 7.43 ± 3.11 µV, p = 0.0021, paired t-test; Fig. 3C).

### Effect of infrared stimulation on the amplitude of local field potentials

Changes in the amplitude of slow waves in the LFP signal typically indicate alterations in neural synchronization, with higher amplitudes reflecting greater network-level synchrony of underlying neural activity (Bernardi et al. 2018; Esser et al. 2007; Riedner et al. 2007; Vyazovskiy et al. 2007). Given that the increase in MUA amplitude during NIR stimulation suggests enhanced neural synchrony, we next asked whether this effect was also reflected in the amplitude of local field potentials. To assess whether local cortical warming via NIR stimulation modulates the temporal synchrony of spontaneous activity, we calculated mean absolute LFP amplitudes during both the stimulation and recovery periods in supragranular (layer 2/3) and infragranular (layer 5) layers of the neocortex.

Absolute LFP amplitudes were significantly smaller in layer 2/3 than in layer 5 (layer 2/3 vs. layer 5; 232.31 ± 71.71 µV vs. 342.85 ± 105.47 µV, p = 0.016, paired t-test). Furthermore, LFP amplitudes in S1Tr were higher compared to PtA, although these differences were not statistically significant, only close to the significance level in layer 5 (PtA vs. S1Tr; layer 2/3: 193.42 ± 53.10 µV vs. 271.19 ± 71.74 µV; p = 0.132, Student’s t-test; layer 5: 276.49 ± 91.69 µV vs. 409.21 ± 76.21 µV, p = 0.0676, Student’s t-test).

About half of the animals showed an increase in the absolute LFP amplitude during stimulation (Fig. 4A), while the other half exhibited a decrease. Thus, when data from all animals were compared, there was no significant change in the LFP amplitudes during stimulation relative to baseline (BL vs. ON; layer 2/3: 232.31 ± 71.71 µV vs. 219.63 ± 50.31 µV; p = 0.4995, paired t-test; layer 5: 342.85 ± 105.47 µV vs. 365.00 ± 92.13 µV, p = 0.446, paired t-test; Fig. 4B).

However, when separating the data based on the cortical location of the optrode, we found region-specific differences. In PtA, the amplitudes of slow waves increased in both layers during stimulation, with a significant effect in layer 5 (BL vs. ON; layer 2/3: 193.42 ± 53.10 µV vs. 218.42 ± 47.75 µV; p = 0.1968, paired t-test; layer 5: 276.49 ± 91.69 µV vs. 353.53 ± 98.78 µV, p = 0.0241, paired t-test; Fig. 4C). In contrast, absolute LFP amplitudes in S1Tr decreased during stimulation in both cortical layers, although the changes did not reach significance (BL vs. ON; layer 2/3: 271.19 ± 71.74 µV vs. 220.83 ± 60.18 µV; p = 0.0636, paired t-test; layer 5: 409.21 ± 76.21 µV vs. 376.47 ± 98.48 µV, p = 0.4096, paired t-test; Fig. 4D).

### Spectral changes in local field potentials during infrared stimulation

To further examine the effects of NIR stimulation on cortical local field potentials, we analyzed spectral changes in the low-frequency band (0.5 – 4 Hz) corresponding to ketamine/xylazine-induced slow waves (0.5 – 2 Hz; Fiáth et al. 2016; Sharma et al. 2010), and high delta activity (2 – 4 Hz) which represents faster delta oscillations. NIR stimulation, accompanied by local cortical warming, induced notable alterations in the spectral amplitude of low-frequency components (Fig. 5A and B).

Similarly to the results of the LFP amplitude analysis, a subset of animals showed a robust increase in the power of the slow wave band (Fig. 5A, left), while others exhibited a marked decrease during stimulation (Fig. 5A, right). These stimulation-induced spectral changes in the LFP reversed rapidly during the recovery period following the cessation of stimulation (Fig. 5A and B). Trial-to-trial variability in spectral changes was relatively low for most animals (e.g., see Fig. 5A).

To quantify these effects, we calculated the spectral amplitude in the two frequency bands described above (i.e., slow wave and high delta bands) for each animal using the last minute of the stimulation (ON) and the recovery periods (BL, Fig. 5C). Spectral amplitudes measured during baseline periods were significantly higher in layer 5 than in layer 2/3 for both frequency bands (layer 2/3 vs. layer 5; slow wave band: 2228.82 ± 691.41 µV vs. 3352.21 ± 1010.75 µV, p = 0.0158, paired t-test; high delta band: 1785.05 ± 765.52 µV vs. 2380.35 ± 1025.41 µV, p = 0.0435, paired t-test). Amplitudes in both the slow wave and high delta bands across layer 2/3 and layer 5 were higher in S1Tr compared to PtA (Table 1), but these differences did not reach statistical significance.

**Table 1.** Spectral amplitudes (in µV) in the two cortical regions (mean ± standard deviation) in the slow wave (0.5 – 2 Hz) and high delta (2 – 4 Hz) frequency bands, and the p-values of Student’s t-tests. PtA, parietal association cortex; S1Tr, trunk region of the primary somatosensory cortex.

|  | Layer | S1Tr (n=4) | PtA (n=4) | p-value |
| --- | --- | --- | --- | --- |
| Slow wave | L2/3 | 2602.79 ± 605.90 | 1854.85 ± 435.60 | 0.133 |
| | L5 | 3907.26 ± 622.82 | 2797.17 ± 885.31 | 0.126 |
| High delta | L2/3 | 2169.58 ± 714.08 | 1400.53 ± 468.94 | 0.170 |
| | L5 | 2958.92 ± 901.00 | 1801.79 ± 599.00 | 0.113 |

When amplitude values from all animals were pooled, there was no difference between stimulation and baseline in either cortical layer or frequency band (Table 2 and 3, Fig. 5C and D). However, when the two cortical areas were analyzed separately, an increase in the average slow wave and high delta amplitudes was found during stimulation in PtA across both cortical layers (Table 2 and 3, Fig. 5E and F). Furthermore, the spectral amplitude of the slow waves was significantly higher in layer 5 during stimulation compared to baseline (Table 2, Fig. 5E). In contrast, in S1Tr the amplitude tended to decrease in both layers and frequency bands, although these differences were not significant (Table 2 and 3, Fig. 5G and H).

**Table 2.** Spectral amplitudes (in µV) in the slow wave (0.5 – 2 Hz) band (mean ± standard deviation), and the p-values of paired t-tests. BL, baseline period; ON, infrared stimulation period; PtA, parietal association cortex; S1Tr, trunk region of the primary somatosensory cortex.

|  | Layer | BL | ON | p-value |
| --- | --- | --- | --- | --- |
| All (n=8) | L2/3 | 2228.82 ± 691.41 | 2098.44 ± 539.63 | 0.413 |
| | L5 | 3352.21 ± 1010.75 | 3495.13 ± 971.30 | 0.517 |
| PtA (n=4) | L2/3 | 1854.85 ± 435.60 | 2040.34 ± 396.02 | 0.246 |
| | L5 | 2797.17 ± 885.31 | 3353.38 ± 797.43 | 0.018 |
| S1Tr (n=4) | L2/3 | 2602.79 ± 605.90 | 2156.55 ± 588.24 | 0.056 |
| | L5 | 3907.26 ± 622.82 | 3636.87 ± 987.38 | 0.401 |

**Table 3.** Spectral amplitudes (in µV) in the high delta (2 – 4 Hz) band (mean ± standard deviation), and the p-values of paired t-tests. BL, baseline period; ON, infrared stimulation period; PtA, parietal association cortex; S1Tr, trunk region of the primary somatosensory cortex.

|  | Layer | BL | ON | p-value |
| --- | --- | --- | --- | --- |
| All (n=8) | L2/3 | 1785.05 ± 765.52 | 1598.34 ± 371.80 | 0.295 |
| | L5 | 2380.35 ± 1025.41 | 2227.93 ± 717.10 | 0.414 |
| PtA (n=4) | L2/3 | 1400.53 ± 468.94 | 1466.69 ± 219.54 | 0.699 |
| | L5 | 1801.79 ± 599.00 | 1871.35 ± 548.00 | 0.284 |
| S1Tr (n=4) | L2/3 | 2169.58 ± 714.08 | 1729.00 ± 398.81 | 0.171 |
| | L5 | 2958.92 ± 901.00 | 2584.52 ± 587.35 | 0.338 |

## Discussion

Slow-wave activity reflects the synchronized behavior of large neuronal populations within the thalamocortical network and exhibits complex spatiotemporal dynamics. In this study, we modulated spontaneously occurring cortical slow-wave activity using near-infrared stimulation in rats anesthetized with ketamine/xylazine. We observed significant changes in multiple properties of slow waves during stimulation, including their amplitudes, the duration of active and silent states and the synchronization level of population spiking activity.

Interestingly, infrared stimulation-induced changes in the LFP (particularly spectral amplitude and slow wave amplitude) varied across the two investigated cortical areas. In the parietal association cortex, NIR neuromodulation enhanced slow and delta oscillations, manifesting as increased spectral amplitude and larger slow wave amplitudes. In contrast, the primary somatosensory cortex showed a reduction in these measures. Several factors may contribute to this interareal discrepancy.

First, anatomical differences between cortical areas may play a role. For instance, the association cortex lacks a well-defined layer 4 and has a reduced overall cortical thickness compared to primary sensory regions. Consequently, although the optrode tip (i.e., the site of stimulation) was consistently positioned 1.2 mm below the cortical surface, its laminar location varied between cortical regions. Based on histological analysis, in PtA, the tip was typically located in the upper to middle part of layer 6, while in S1Tr it was in the lower part of layer 5. These laminar differences may have affected the spatial distribution and intensity of temperature changes induced by infrared stimulation.

Second, the size of the craniotomy may have influenced the baseline temperature of the cortical tissue near the optrode. In PtA recordings, the craniotomy (10 x 5 mm^2^) was more than five times larger than in S1Tr recordings (3 x 3 mm^2^). Prior work has demonstrated that exposed cortical tissue in a craniotomy is significantly cooler than physiological brain temperature due to heat loss (Kalmbach and Waters 2012). In mice, the exposed cortical surface can be up to 10 °C below core body temperature, markedly affecting ongoing cortical activity (Kalmbach and Waters 2012). A larger craniotomy would thus produce greater heat dissipation, possibly lowering baseline cortical temperature more and perturbing baseline neural activity at a higher degree and on a larger spatial scale. This factor may partially explain the region-specific effects of IR stimulation observed here.

The observed increase in slow/delta amplitude in PtA is consistent with previous findings. For instance, in ferret coronal brain slices from the primary visual cortex expressing slow rhythms in vitro, higher chamber temperatures were associated with increased spectral power in low LFP frequencies (Reig et al. 2010). Similarly, Sheroziya and Timofeev (2015) reported elevated delta band (0.2 - 4 Hz) power in the motor cortex of head-fixed, non-anesthetized mice at physiological cortical temperatures compared to cooled tissue. Interestingly, contrary to our results in S1Tr, in ketamine/xylazine anesthetized mice, the same study found that higher tissue temperatures in the somatosensory cortex were associated with larger slow wave amplitudes (Sheroziya and Timofeev 2015). Kalmbach and Waters (2012) also reported increased power in the 0–1.5 Hz range at physiological brain temperatures in the barrel cortex of mice anesthetized with isoflurane compared to those measured in cranial windows with lower cortical temperatures.

In contrast to the regional variability of LFP-related measures, other parameters, namely state durations and MUA amplitude, showed consistent changes across animals. Specifically, infrared stimulation shortened up-states and prolonged down-states, accompanied by increased MUA amplitudes and steeper MUA slopes in both supragranular and infragranular layers.

Our findings regarding state durations align partially with above studies. Higher cortical temperatures have been associated with shorter up-states in both the barrel cortex of anesthetized mice and visual cortical slices (Kalmbach and Waters 2012; Reig et al. 2010). However, the prolongation of down-states observed in our study during infrared stimulation was only reported by Kalmbach and Waters (2012). In brain slices, the duration of down-states initially decreased with temperature increases from 32 °C to 36 °C, but then began to lengthen above 38 °C.

In anesthetized rats, infrared stimulation also produced a steeper rise in cortical MUA and higher peak amplitudes during up-states. These findings are consistent with those from brain slice experiments, where warmer chamber temperatures led to sharper population activity during state transitions (Reig et al. 2010), potentially reflecting more synchronized neuronal firing with a faster recruitment of local neuronal populations.

Our previous study using a similar optrode device showed that infrared stimulation enhanced low-frequency (1-3 Hz) LFP power and increased MUA amplitude in the neocortex of anesthetized rats (Horváth et al. 2024a). The optrode in this report featured a micromirror tip that allowed for spatially focused delivery of infrared light focused to one side of the silicon shaft. The study found significantly higher delta power and MUA amplitude within 300 μm in front of the optrode shaft, where heating was maximal, compared to the rear side of the device (Horváth et al. 2024a). In some of these experiments, decreased spectral and MUA amplitudes were observed in backside regions less affected by infrared stimulation compared to baseline values. These findings suggest that the spatial reach of infrared stimulation-induced neuromodulation is relatively localized, likely limited to a few hundred micrometers from the optrode tip.

Since the optrode applied in this study contained relatively few and relatively large microelectrodes, yielding only a small number of well-isolated single-units, single-unit analysis was not performed here. Nevertheless, recent studies in rats have investigated the influence of infrared stimulation at the level of single neurons, reporting both excitatory and suppressive effects on firing rates (Balogh-Lantos et al. 2024; Horváth et al. 2020; 2022; 2024a). Under continuous-wave stimulation, as applied here, roughly one-third of identified units exhibited reduced firing rates (∼45% decrease on average compared to baseline), while approximately one-quarter showed increased firing (∼110% mean increase in firing rates) in anesthetized rats (Balogh-Lantos et al. 2024). Despite this bidirectional modulation, the overall effect of these changes was a modest net increase in neuronal firing, which likely accounts for the enhanced population activity (MUA) observed in our study. Moreover, because infrared stimulation shortened up-states, spikes became confined to briefer periods, producing sharper, higher-amplitude MUA peaks with steeper slopes. This finding further supports the conclusion that infrared stimulation promotes more synchronized slow-wave activity.

In this study, we focused on the effects of infrared stimulation on the activity of neural circuits. However, other cortical cell types, particularly glial cells, may also contribute to the observed changes. For instance, calcium imaging combined with LFP recordings has shown that astrocytes play a role in the generation of slow-wave activity in the primary visual cortex of ketamine/xylazine anesthetized rats (Szabó et al. 2017). Furthermore, calcium imaging in the somatosensory cortex of urethane-anesthetized rats revealed that pulsed infrared light elicits both slow and fast responses, with the slow component originating from astrocytes (Cayce et al. 2014). More recently, a comprehensive study using calcium imaging, pharmacology, electrophysiology, and genetic manipulation in rat cortical astrocyte cultures provided detailed insight into how pulsed infrared light modulates astrocytic activity (Borrachero-Conejo et al. 2020). Taken together, these findings suggest that astrocytes may also contribute to the NIR stimulation-related modulation of cortical slow wave dynamics observed in our study.

## Acknowledgements

This work is dedicated to the memory of our coauthor Zoltán Fekete (deceased December 2024), whose contributions were essential to this study.

## Author contributions

Ágnes Szabó (Data curation, Formal analysis, Methodology, Software, Validation, Visualization, Writing— original draft, Writing—review & editing), Richárd Fiáth (Conceptualization, Data curation, Investigation, Methodology, Resources, Supervision, Funding acquisition, Writing— original draft, Writing—review & editing), Ágoston Csaba Horváth (Data curation, Investigation, Writing—review & editing), Péter Barthó (Formal analysis, Writing—review & editing), István Ulbert (Funding acquisition, Supervision, Writing—review & editing) and Zoltán Fekete (Conceptualization, Methodology, Project administration, Supervision, Funding acquisition, Writing— original draft).

## References

Achermann P, Borbély AA. 1997. Low-frequency (< 1 Hz) oscillations in the human sleep electroencephalogram. Neuroscience. 81:213–222. 10.1016/s0306-4522(97)00186-3.

Balogh-Lantos Z, Fiáth R, Horváth ÁC, Fekete Z. 2024. High density laminar recordings reveal cell type and layer specific responses to infrared neural stimulation in the rat neocortex. Sci Rep. 14:31523. 10.1038/s41598-024-82980-w.

Barth AL, Poulet JF. Experimental evidence for sparse firing in the neocortex. 2012. Trends Neurosci. 35:345–355. 10.1016/j.tins.2012.03.008.

Beltramo R et al. 2013. Layer-specific excitatory circuits differentially control recurrent network dynamics in the neocortex. Nat Neurosci. 16:227–234. 10.1038/nn.3306.

Bernardi G, Siclari F, Handjaras G, Riedner BA, Tononi G. 2018. Local and widespread slow waves in stable NREM sleep: evidence for distinct regulation mechanisms. Front Hum Neurosci. 12:248. 10.3389/fnhum.2018.00248.

Bokil H, Andrews P, Kulkarni JE, Mehta S, Mitra PP. 2010. Chronux: a platform for analyzing neural signals. J Neurosci Methods. 192:146–151. 10.1016/j.jneumeth.2010.06.020.

Born J, Rasch B, Gais S. 2006. Sleep to remember. Neuroscientist. 12:410–424. 10.1177/1073858406292647.

Borrachero-Conejo AI, et al. 2020. Stimulation of water and calcium dynamics in astrocytes with pulsed infrared light. FASEB J. 34:6539–6553. 10.1096/fj.201903049R.

Cayce JM, Friedman RM, Jansen ED, Mahavaden-Jansen A, Roe AW. 2011. Pulsed infrared light alters neural activity in rat somatosensory cortex in vivo. Neuroimage. 57:155–166. 10.1016/j.neuroimage.2011.03.084.

Cayce, JM et al. 2014. Calcium imaging of infrared-stimulated activity in rodent brain. Cell Calcium. 55:183–190. 10.1016/j.ceca.2014.01.004.

Chauvette S, Volgushev M, Timofeev I. 2010. Origin of active states in local neocortical networks during slow sleep oscillation. Cereb Cortex. 20:2660–2674. 10.1093/cercor/bhq009.

Clement EA et al. 2008. Cyclic and sleep-like spontaneous alternations of brain state under urethane anaesthesia. PloS One. 3:e2004. 10.1371/journal.pone.0002004.

Crunelli V, Hughes SW. 2010. The slow (< 1 Hz) rhythm of non-REM sleep: a dialogue between three cardinal oscillators. Nat Neurosci. 13:9–17. 10.1038/nn.2445.

Csernyus B et al. 2021. A multimodal, implantable sensor array and measurement system to investigate the suppression of focal epileptic seizure using hypothermia. J Neural Eng, 18:0460c3. 10.1088/1741-2552/ac15e6.

D’Andola M et al. 2018. Bistability, causality, and complexity in cortical networks: an in vitro perturbational study. Cereb Cortex. 28:2233–2242. 10.1093/cercor/bhx122.

Dasilva M et al. 2021. Modulation of cortical slow oscillations and complexity across anesthesia levels. Neuroimage. 224:117415. 10.1016/j.neuroimage.2020.117415.

Esser SK, Hill SL, Tononi G. 2007. Sleep homeostasis and cortical synchronization: I. Modeling the effects of synaptic strength on sleep slow waves. Sleep. 30:1617–1630. 10.1093/sleep/30.12.1617.

Fiáth R et al. 2016. Laminar analysis of the slow wave activity in the somatosensory cortex of anesthetized rats. Eur J Neurosci. 44:1935–1951. 10.1111/ejn.13274.

Fiáth, R. et al. 2019. Slow insertion of silicon probes improves the quality of acute neuronal recordings. Sci Rep. 9:111. 10.1038/s41598-018-36816-z.

Fekete Z, Horváth ÁC, Zátonyi A. 2020. Infrared neuromodulation: a neuroengineering perspective. J Neural Eng. 17:051003. 10.1088/1741-2552/abb3b2.

Funk CM, Honjoh S, Rodriguez AV, Cirelli C, Tononi G. 2016. Local slow waves in superficial layers of primary cortical areas during REM sleep. Curr Biol. 26:396–403. 10.1016/j.cub.2015.11.062.

Greenberg A, Dickson CT. 2013. Spontaneous and electrically modulated spatiotemporal dynamics of the neocortical slow oscillation and associated local fast activity. Neuroimage. 83:782–794. 10.1016/j.neuroimage.2013.07.034.

Hablitz LM et al. 2019. Increased glymphatic influx is correlated with high EEG delta power and low heart rate in mice under anesthesia. Sci Adv. 5:eaav5447. 10.1126/sciadv.aav5447.

Hangya B et al. 2011. Complex propagation patterns characterize human cortical activity during slow-wave sleep. J Neurosci, 31:8770–8779. 10.1523/JNEUROSCI.1498-11.2011.

Hasenstaub A, Sachdev RN, McCormick DA. 2007. State changes rapidly modulate cortical neuronal responsiveness. J Neurosci. 27:9607–9622. 10.1523/JNEUROSCI.2184-07.2007.

Horváth ÁC et al. 2018. A multimodal microtool for spatially controlled infrared neural stimulation in the deep brain tissue. Sens Actuators B Chem. 263:77–86. 10.1016/j.snb.2018.02.034.

Horváth ÁC et al. 2020. Infrared neural stimulation and inhibition using an implantable silicon photonic microdevice. Microsyst Nanoeng. 6:44. 10.1038/s41378-020-0153-3.

Horváth ÁC et al. 2022. Histological and electrophysiological evidence on the safe operation of a sharp-tip multimodal optrode during infrared neuromodulation of the rat cortex. Sci Rep. 12:11434. 10.1038/s41598-022-15367-4.

Horváth ÁC et al. 2024a. Silicon Optrode with a Micromirror-Tip Providing Tunable Beam Profile During Infrared Neuromodulation of the Rat Neocortex. Adv Mater Technol. 9:2400044. 10.1002/admt.202400044.

Horváth C, Ulbert I, Fiáth R. 2024b. Propagating population activity patterns during spontaneous slow waves in the thalamus of rodents. NeuroImage. 285:120484. 10.1016/j.neuroimage.2023.120484.

Ismaiel E, Fiáth R, Szabó Á, Horváth ÁC, Fekete Z. 2023. Thermal neuromodulation using pulsed and continuous infrared illumination in a penicillin-induced acute epilepsy model. Sci Rep. 13:14460. 10.1038/s41598-023-41552-0.

Kalmbach AS, Waters J. 2012. Brain surface temperature under a craniotomy. J Neurophysiol. 108:3138–3146. 10.1152/jn.00557.2012.

Kuki T et al. 2013. Frequency-dependent entrainment of neocortical slow oscillation to repeated optogenetic stimulation in the anesthetized rat. Neurosci Res. 75:35–45. 10.1016/j.neures.2012.10.007.

Lőrincz M et al. 2015. A distinct class of slow (∼ 0.2–2 Hz) intrinsically bursting layer 5 pyramidal neurons determines UP/DOWN state dynamics in the neocortex. J Neurosci. 35:5442–5458. 10.1523/JNEUROSCI.3603-14.2015.

Massimini M, Huber R, Ferrarelli F, Hill S, Tononi G. 2004. The sleep slow oscillation as a traveling wave. J Neurosci. 24:6862–6870. 10.1523/JNEUROSCI.1318-04.2004.

Massimini M et al. 2007. Triggering sleep slow waves by transcranial magnetic stimulation. Proc Natl Acad Sci. 104:8496–8501. 10.1073/pnas.0702495104.

Neske GT. 2016. The slow oscillation in cortical and thalamic networks: mechanisms and functions. Front Neural Circuits. 9:88. 10.3389/fncir.2015.00088.

Nir Y et al. 2011. Regional slow waves and spindles in human sleep. Neuron. 70:153–169. 10.1016/j.neuron.2011.02.043.

Paxinos G, Watson C. 2006. The rat brain in stereotaxic coordinates: hard cover edition. Elsevier.

Rasch B, Born J. 2013. About sleep’s role in memory. Physiol Rev. 93:681–766. 10.1152/physrev.00032.2012.

Reig R, Mattia M, Compte A, Belmonte C, Sanchez-Vives MV. 2010. Temperature modulation of slow and fast cortical rhythms. J Neurophysiol. 103:1253–1261. 10.1152/jn.00890.2009.

Riedner BA et al. 2007. Sleep homeostasis and cortical synchronization: III. A high-density EEG study of sleep slow waves in humans. Sleep. 30:1643–1657. 10.1093/sleep/30.12.1643.

Sakata S, Harris KD. 2009. Laminar structure of spontaneous and sensory-evoked population activity in auditory cortex. Neuron. 64:404–418. 10.1016/j.neuron.2009.09.020.

Sanchez-Vives MV, McCormick DA. 2000. Cellular and network mechanisms of rhythmic recurrent activity in neocortex. Nat Neurosci. 3:1027–1034. 10.1038/79848.

Sharma AV, Wolansky T, Dickson CT. 2010. A comparison of sleeplike slow oscillations in the hippocampus under ketamine and urethane anesthesia. J Neurophysiol. 104:932–939. 10.1152/jn.01065.2009.

Sheroziya M, Timofeev I. 2014. Global intracellular slow-wave dynamics of the thalamocortical system. J Neurosci. 34:8875–8893. 10.1523/JNEUROSCI.4460-13.2014.

Sheroziya M, Timofeev I. 2015. Moderate cortical cooling eliminates thalamocortical silent states during slow oscillation. J Neurosci. 35:13006–13019. 10.1523/JNEUROSCI.1359-15.2015.

Shu Y, Hasenstaub A, McCormick DA. 2003. Turning on and off recurrent balanced cortical activity. Nature. 423:288–293. 10.1038/nature01616.

Siclari F, Tononi G. 2017. Local aspects of sleep and wakefulness. Curr Opin Neurobiol. 44:222–227. 10.1016/j.conb.2017.05.008.

Steriade M, Nunez A, Amzica F. 1993. A novel slow (< 1 Hz) oscillation of neocortical neurons in vivo: depolarizing and hyperpolarizing components. J Neurosci. 13:3252–3265. 10.1523/JNEUROSCI.13-08-03252.1993.

Stroh A et al. 2013. Making waves: initiation and propagation of corticothalamic Ca2+ waves in vivo. Neuron, 77:1136–1150. 10.1016/j.neuron.2013.01.031.

Szabó Z et al. 2017. Extensive astrocyte synchronization advances neuronal coupling in slow wave activity in vivo. Sci Rep. 7:6018. 10.1038/s41598-017-06073-7.

Timofeev I, Grenier F, Bazhenov M, Sejnowski TJ, Steriade M. 2000. Origin of slow cortical oscillations in deafferented cortical slabs. Cereb Cortex. 10:1185–1199. 10.1093/cercor/10.12.1185.

Timofeev I, Chauvette S. 2017. Sleep slow oscillation and plasticity. Curr Opin Neurobiol. 44:116–126. 10.1016/j.conb.2017.03.019.

Tononi G, Cirelli C. 2006. Sleep function and synaptic homeostasis. Sleep Med Rev. 10:49–62. 10.1016/j.smrv.2005.05.002.

Tononi G, Cirelli C. 2014. Sleep and the price of plasticity: from synaptic and cellular homeostasis to memory consolidation and integration. Neuron. 81:12–34. 10.1016/j.neuron.2013.12.025.

Vyazovskiy VV, Riedner BA, Cirelli C, Tononi G. 2007. Sleep homeostasis and cortical synchronization: II. A local field potential study of sleep slow waves in the rat. Sleep. 30:1631–1642. 10.1093/sleep/30.12.1631.

Vyazovskiy VV, Faraguna U, Cirelli C, Tononi G. 2009. Triggering slow waves during NREM sleep in the rat by intracortical electrical stimulation: effects of sleep/wake history and background activity. J Neurophysiol. 101:1921–1931. 10.1152/jn.91157.2008.

Vyazovskiy VV et al. 2011. Local sleep in awake rats. Nature. 472:443–447. 10.1038/nature10009.

Wang X et al. 2019. Transcranial photobiomodulation with 1064-nm laser modulates brain electroencephalogram rhythms. Neurophotonics. 6:025013–025013. 10.1117/1.NPh.6.2.025013.

Ward-Flanagan R, Lo AS, Clement EA, Dickson CT. 2022. A comparison of brain-state dynamics across common anesthetic agents in male Sprague-Dawley rats. Int J Mol Sci. 23:3608. 10.3390/ijms23073608.

Wester JC, Contreras D. 2012. Columnar interactions determine horizontal propagation of recurrent network activity in neocortex. J Neurosci. 32:5454–5471. 10.1523/JNEUROSCI.5006-11.2012.

Xie L et al. 2013. Sleep drives metabolite clearance from the adult brain. Science. 342:373–377. 10.1126/science.1241224.

Zomorrodi R, Loheswaran G, Pushparaj A, Lim L. 2019. Pulsed near infrared transcranial and intranasal photobiomodulation significantly modulates neural oscillations: a pilot exploratory study. Sci Rep. 9:6309. 10.1038/s41598-019-42693-x.

